# miR-29a-3p and TGF-β Axis in Fanconi Anemia: Mechanisms Driving Metabolic Dysfunction and Genome Stability

**DOI:** 10.1101/2025.01.14.632746

**Authors:** Nadia Bertola, Stefano Regis, Vanessa Cossu, Matilde Balbi, Martina Serra, Fabio Corsolini, Cristina Bottino, Paolo Degan, Carlo Dufour, Filomena Pierri, Enrico Cappelli, Silvia Ravera

## Abstract

Fanconi anemia (FA) is a rare genetic disorder characterized by bone marrow failure and cancer susceptibility due to defective DNA double-strand break repair. However, FA cells are also characterized by mitochondrial dysfunction and redox imbalance. To identify a common factor among these alterations, we focused on miR-29a-3p, a microRNA involved in hematopoiesis. Our data show that miR-29a-3p is downregulated in lymphoblasts and fibroblasts mutated for the FANC-A gene, causing the overexpression of its target genes, FOXO3, SGK1, and IGF1, which results in PI3K/AKT pathway hyperactivation, altered mitochondrial metabolism and insufficient antioxidant response. Furthermore, miR-29a-3p downregulation appears associated with hyperactivation of the TGF-β signal. However, restoring miR-29a-3p expression improves mitochondrial metabolism, oxidative stress response, and DNA damage repair by inhibiting the PI3K/AKT pathway and modulating TGF-β signaling by a feedback mechanism. Based on these findings, miR-29a-3p appears as a promising molecular target to address several mechanisms based on FA pathogenesis.

## INTRODUCTION

Fanconi anemia (FA) is a genetic disease, typically with pediatric onset, characterized by aplastic anemia and predisposition to several types of cancer, such as acute myeloid leukemia, gynecological carcinoma, and head and neck squamous cell carcinoma (HNSCC) ^1,2^.

DNA repair has long been considered the primary defect in FA cells ^3,4^. However, in the last 20 years, research has shown that FA proteins are involved in several other cellular processes, including defective energy metabolism ^5,6^, impaired antioxidant response^7–9^, and the overproduction of pro-inflammatory and cytotoxic cytokines ^10–13^. In detail, FA cells are characterized by altered electron transport between respiratory complexes I and III ^14,15^, leading to the uncoupling of oxidative phosphorylation (OxPhos) ^16–18^. As a result, FA cells exhibit increased reactive oxygen species (ROS) production ^19,20^, not counteracted by endogenous antioxidant defensesbecause FA cells are not able to trigger an adaptative response to the oxidative stress increment ^21–23^. This imbalance in redox status promotes DNA damage accumulation ^24^ and the development of a pro-inflammatory environment ^7^, triggering a vicious circle. Additionally, unrepaired DNA damage is reported to contribute to the pro-inflammatory conditions observed in FA patients ^25,26^, increasing the expression of cytotoxic cytokines. This heightened expression is associated with increased cellular sensitivity and leads to the death of hematopoietic progenitors ^13,27^.In this context, several pro-inflammatory cytokines have already been associated with defective hematopoiesis in patients with FA ^28^. For example, bone marrow mononuclear cells (BM-MNC) isolated from FA patients have shown hypersensitivity to TNF-α, which inhibits erythropoiesis *in vitro*^12^. Additionally, hyperactivity of the TGF-β/SMAD3 pathway inhibits DNA repair via homologous recombination (HR), activating non-homologous end joining (NHEJ), which is toxic to FA HSCs ^29^. Although DNA damage accumulation, impaired energy metabolism and antioxidant defenses, and increased production of proinflammatory cytokines in FA have been extensively described in the literature, no mechanism has yet been proposed yet to explain their interconnection and mutual influence. To address this gap, we focused our attention on the microRNA (miRNA) profile in FA cells, because, to date, few studies have explored the miRNA profile in FA cells ^30,31^. In this study, we investigate the role of miR-29a-3p, a member of the miR-29 family, which modulates numerous biochemical and physiological pathways, including the maturation, differentiation, and survival of hematopoietic stem and progenitor cells (HSPCs) ^32^. Notably, miR-29a-3p has been shown to regulate DNA methyltransferase 3 (DNMT3), supporting self-renewal in HSPCs and guiding lineage commitment and differentiation ^32,33^.

Our findings reveal that FA lymphoblasts and primary fibroblasts exhibit significantly reduced expression of miR-29a-3p compared to healthy controls and isogenic corrected cells. This reduction is mediated by negative feedback involving TGF-β pathway activity. Furthermore, transfection of FA cells with miR-29a-3p restores mitochondrial function, enhances the oxidative stress response and reduces DNA damage by modulating the TGF-β pathway through decreasingSMAD3phosphorylation.

## RESULTS

### miR-29a-3p expression is downregulated in Fanc-A cells

Although several miRNAs may potentially be involved in the pathogenesis of FA^30,31^,we focused our attention on miR-29a-3p, due to its role in modulating mitochondrial activity and redox balance^32^, two features altered in FA cells ^17,21^. Our data show a significant reduction in miR-29a-3p expression in Fanc-A lymphoblasts(Figure 1A) and fibroblasts (Figure S1) compared to the corresponding isogenic corrected cells, suggesting a possible role of miR-29a-3p in mitochondrial dysfunction and associated oxidative stress production in FA cells.

**Figure 1.**
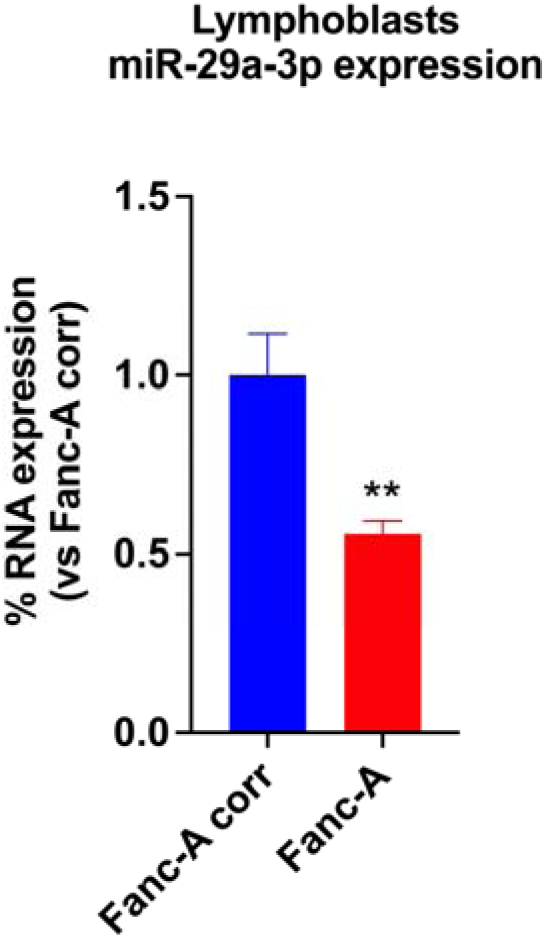
miR-29a-3p expression in Fanc-A cells. The graph shows the comparison of miR-29a-3p expression in isogenic Fanc-A corrected lymphoblasts (Fanc-A corr, used as a control) and Fanc-A lymphoblasts (Fanc-A). RNU44 was used as a reference control. Data are expressed as mean ± SD and are representative of three independent experiments. ** indicates a significant difference for p < 0.01 between Fanc-A corr and Fanc-A.

### Fanc-A cells display an alteration of several putative miR-29a-3p-regulated genes involved in DNA repair, mitochondrial function, redox balance, and apoptosis

Based on the results of miR-29a-3p expression, we interrogated the miRPathDB v2.0 database to identify putative associations between miRNAs, target genes, and cellular pathways. Specifically, we selected genes potentially regulated by miR-29a-3p and involved in DNA damage response, oxidative stress, mitochondrial metabolism, lipid metabolism, and apoptosis, all of which are pathways potentially altered in FA cells. Furthermore, TargetScan was used to define the gene list based on the strength of predicted miRNA-target gene interactions. The list was then refined by analyzing the role and subcellular location of each gene using NCBI Gene and GeneCards. The final list is reported in Table 1.

**Table 1.** Putative target genes of miR-29a-3p in DNA damage response processes, mitochondrial function, oxidative stress, lipid metabolism, and apoptosis.

| *Putative miR-29a-3p-regulated genes* |  |  |  |  |
| --- | --- | --- | --- | --- |
| **DNA damage response** | **Mitochondrion** | **Oxidative stress** | **Lipid metabolism** | **Apoptosis** |
| BTG2 | BAK1 | CCNA2 | CPS1 | BBC3 |
| CTC1 | BBC3 | FOS | **FOXO3** | DIABLO |
| FEM1B | CPS1 | **FOXO3** | FOXP1 | FEM1B |
| **FOXO3** | CYCS | KDM6B | GPAM | **FOXO3** |
| MORF4L1 | DHTKD1 | KLF4 | HMGCS1 | GSK3B |
| MYCN | DIABLO | MMP2 | KLF4 | **IGF1** |
| PIK3R1 | **FOXO3** | REST | LRP6 | ITGA6 |
| POLD3 | GACTC | TNFAIP3 | MAPK8 | MAZ |
| PPM1D | GSK3B | PDGFRB | PDK4 | PIK3R1 |
| RNF138 | HRK | PXDN | PPARD | PPP1R13B |
| **SGK1** | MAPK8 | **IGF1** | PPARGC1A | PTEN |
| TDG | PDHX | **SGK1** | RARB | TNFAIP3 |
| BBC3 | PPARGC1A |  | REST | **IGF1** |
| CCND2 | PTEN |  | RORA |  |
| CDK6 | **SGK1** |  | SIRT1 |  |
|  | SIRT1 |  | THRA |  |
|  | SLC25A15 |  | TNFAIP3 |  |
|  | UBIAD1 |  | TRIM24 |  |
|  | **IGF1** |  |  |  |

### miR-29a-3p transfection reduced the oxidative damage and improved the oxidative phosphorylation in Fanc-A cells

Since low expression of miR-29a-3p in FA cells suggests possible altered expression of genes involved in oxidative stress response (Table 1), catalase (CAT) activity, and intracellular malondialdehyde (MDA) levels have been evaluated as markers of antioxidant defenses and oxidative damage, in FA cells transfected with miR-29a-3p. Additionally, 8-hydroxy-2’-deoxyguanosine (8-OHdG) content and histone H2AX phosphorylation have been assayed as DNA damage markers.

Fanc-A cells displayed reduced CAT activity (Figure 2A and Figure S2A) and increased concentrations of MDA (Figure 2B and Figure S2B) and 8-OHdG (Figure 2C and Figure S2C) compared to the corrected cells, indicating increased oxidative damage to lipids and DNA due to defective antioxidant defenses. In addition, histone H2AX was hyper-phosphorylated in Fanc-A lymphoblasts (Figure 2D), confirming DNA damage accumulation. However, these detrimental effects appeared partially recovered after miR-29a-3p transfection, suggesting a role of this miRNA in the unbalanced oxidative stress of FA cells.

**Figure 2.**
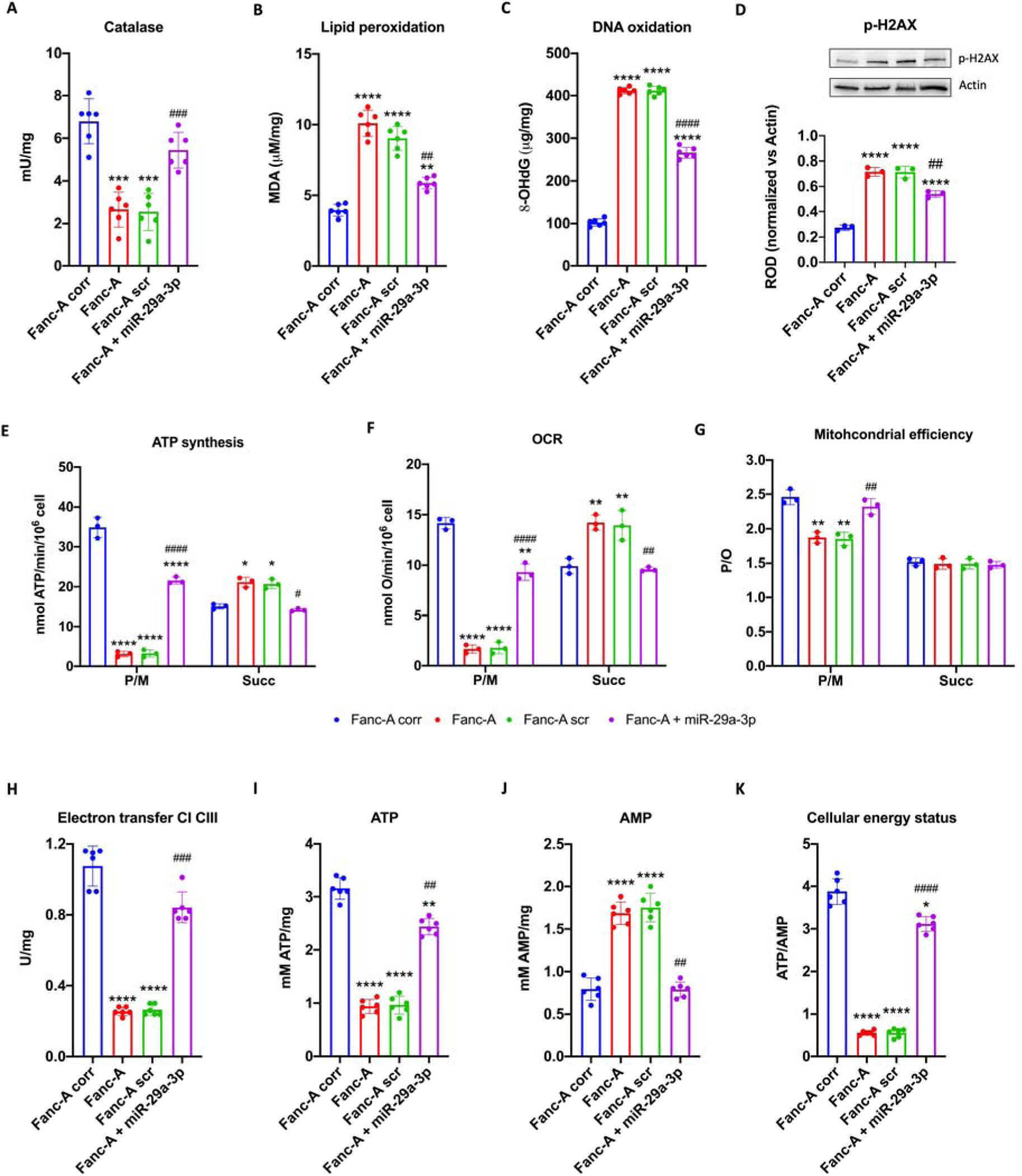
Antioxidant defenses, oxidative stress, and energy metabolism were modulated by miR-29a-3p expression in Fanc-A lymphoblasts. All analyses were conducted on Fanc-A lymphoblasts corrected with the WT Fanc-A gene (Fanc-A corr), Fanc-A lymphoblasts (Fanc-A), Fanc-A lymphoblasts transfected with amiRNA mimic negative control for 48h (Fanc-A scr), and Fanc-A lymphoblasts transfected with miR-29a-3p for 48h (Fanc-A + miR-29a-3p). (A) Catalase activity as an antioxidant defense marker. (B) Intracellular concentration of malondialdehyde (MDA) as a lipid peroxidation marker. (C) 8-hydroxy-2’-deoxyguanosine (8-OHdG) content as a DNA oxidation marker. (D) WB signal and relative densitometric analysis of p-H2AX. The densitometric analysis was normalized to the actin signal and used as a housekeeping protein. (E) ATP synthesis through F_o_F_1_-ATP synthase. (F) Oxygen consumption rate (OCR). (G) P/O value, an OxPhos efficiency marker. For Panels E, F, and G, the analyses were conducted in the presence of pyruvate plus malate (P/M) or succinate (Succ) to induce the OxPhos pathways led by Complex I or Complex II, respectively. (H) Electron transfer between Complexes I and III. (I) Intracellular ATP content. (J) Intracellular AMP content. (K) Cellular energy status is obtained by calculating the ATP/AMP ratio. Data are expressed as mean ± SD and are representative of three independent experiments for Panels E-G and six independent experiments for Panels A-D and H-K. **, ***, and **** indicate a significant difference for p < 0.01, 0.001, or 0.0001, respectively, between Fanc-A corr and Fanc-A or Fanc-A scr. ##, ###, and #### indicate a significant difference for p < 0.01, 0.001, or 0.0001, respectively, between Fanc-A + miR-29a-3p and Fanc-A or Fanc-A scr.

Considering that miR-29a-3p could also regulate genes involved in energy metabolism and that FA cells display an altered OxPhos associated with increased oxidative stress production ^14,17,34^, the ATP synthesis through F_o_F_1_-ATP synthase, the oxygen consumption rate (OCR), and the OxPhos efficiency were investigated before and after miR-29a-3p transfection in Fanc-A lymphoblasts and fibroblasts. As expected, data show that Fanc-A cells were characterized by defective ATP production (Figure 2E and Figure S2D) and OCR (Figure 2F and Figure S2E) when OxPhos was induced by pyruvate/malate addition but not with succinate. This impairment resulted in a decrease of OxPhos efficiency via complexes I-III-IV, as indicated by the reduction of theP/O value (Figure 2G and Figure S2F). All these metabolic parameters showed a significant improvement after miR-29a-3p transfection. This recovery depended on the restoration of electron transport between respiratory complexes I and III (Figure 2H and Figure S2G), leading to an increase in intracellular ATP content (Figure 2I and Figure S2H) and a reduction in AMP concentration (Figure 2J and Figure S2I), ultimately improving the energy status (Figure 2K and Figure S2J).

### miR-29a-3p transfection restores FOXO3, SGK1, and IGF1 genes expression in Fanc-A cells

The data presented in Table 1 highlight that FOXO3, SGK1, and IGF1 are implicated in multiple pathways, including DNA damage response, mitochondrial activity regulation, oxidative stress management, and apoptosis, all hallmark features of FA pathology. Consequently, the expression of these genes was assessed in Fanc-A lymphoblasts (Figure 3) and Fanc-A fibroblasts (Figure S3). The results revealed significantly elevated expression levels for all three genes compared to their corrected counterparts. By contrast, in Fanc-A cells transfected with miR-29a-3p, the expression of the FOXO3, SGK, and IGF1 genes decreased by about 40%, 34%, and 30%, respectively, compared to the FA cells, reaching a comparable level compared to control cells.

**Figure 3.**
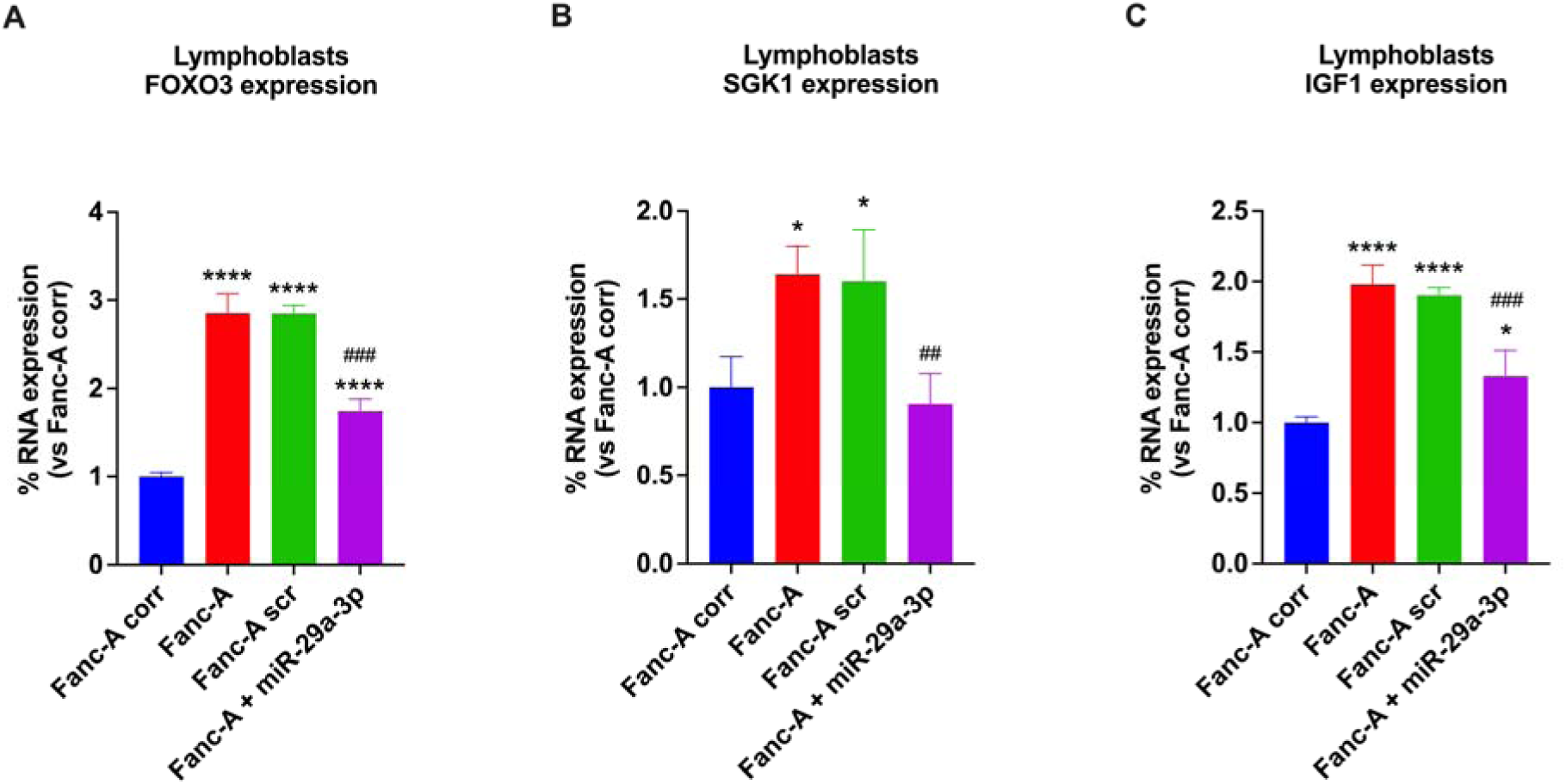
FOXO3, SGK1, and IGF1 expression in Fanc-A lymphoblasts. Graphs show the comparison of FOXO3 (A), SGK1 (B), and IGF1 (C) expression in (i) Fanc-A cells corrected with the WT Fanc-A gene (Fanc-A corr), (ii) Fanc-A cells (Fanc-A), (iii) Fanc-A cells transfected with a miRNA mimic negative control for 48h (Fanc-A scr), and (iv) Fanc-A cells transfected with miR-29a-3p for 48h (Fanc-A + miR-29a-3p). GAPDH was used as the reference control. Data are expressed as mean ± SD and are representative of three independent experiments. * or **** indicate a significant difference for p < 0.05 or 0.0001, respectively, between Fanc-A corr and the other samples. ## and ### indicate a significant difference for p < 0.01 or 0.001, respectively, between Fanc-A + miR-29a-3p and Fanc-A or Fanc-A scr.

### Treatment with miR-29a-3p restores the nuclear translocation of FOXO3a in Fanc-A lymphoblasts

Since FOXO3a function is closely linked to its translocation from the cytoplasm to the nucleus^35^, FOXO3a localization has been evaluated by WB analysis, observing thatFanc-A lymphoblasts predominantly expressed FOXO3a in the cytoplasmic fraction while Fanc-A corr cellsexhibited higher protein expression in the nucleus. Conversely, miR-29a-3p transfection in Fanc-A cells significantly promoted FOXO3a translocation to the nucleus, enhancing its transcriptional activity (Figure 4).

**Figure 4.**
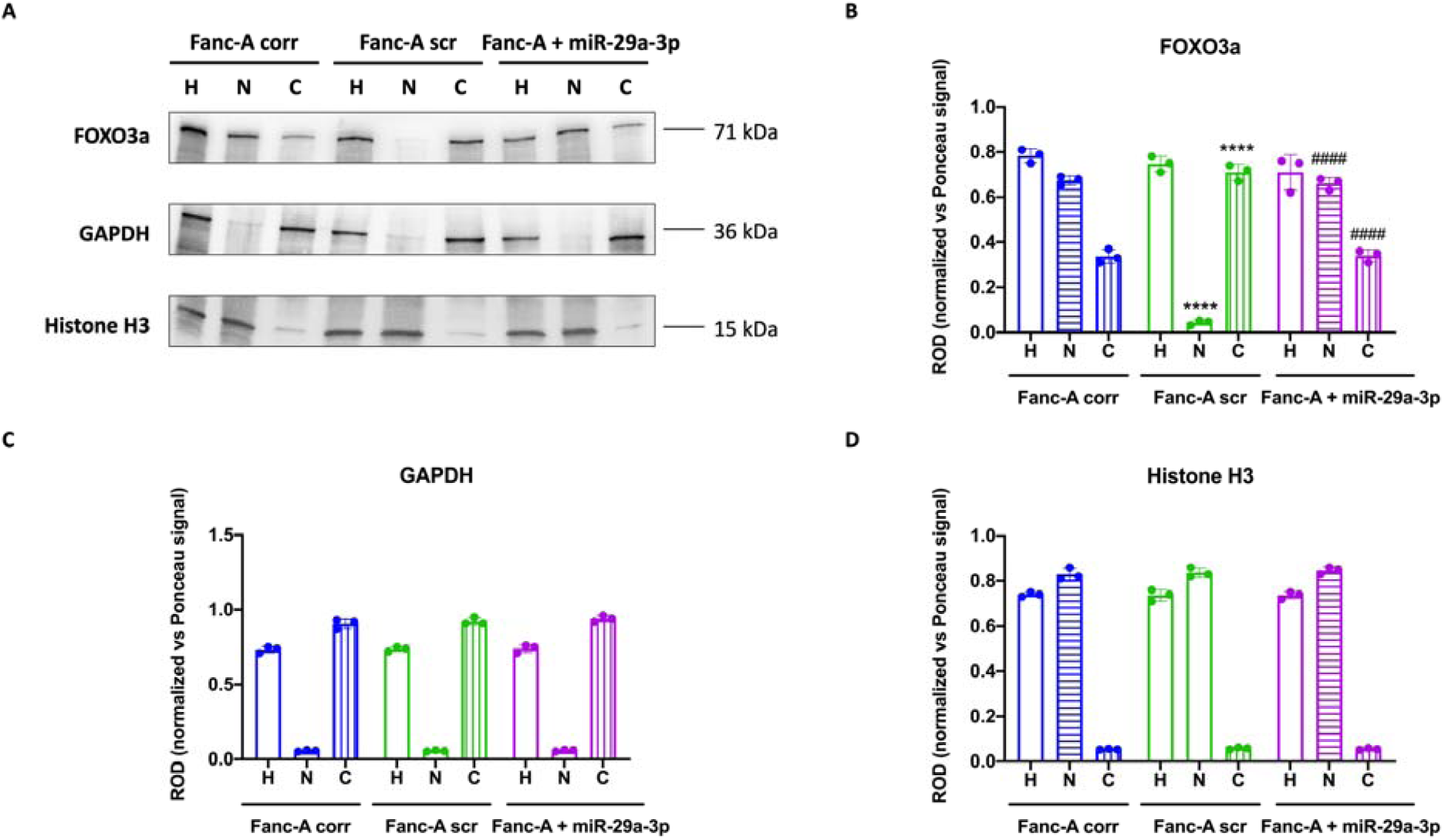
miR-29a-3p changed FOXO3a intracellular localization in Fanc-A lymphoblasts. All analyses were conducted on cell homogenate (H), nuclear fraction (N), and cytoplasmic fraction (C) derived from Fanc-A lymphoblasts corrected with the WT Fanc-A gene (Fanc-A corr), Fanc-A lymphoblasts transfected with a miRNA mimic negative control for 48h (Fanc-A scr), and Fanc-A lymphoblasts transfected with miR-29a-3p for 48h (Fanc-A + miR-29a-3p). (A)Representative WB signals of FOXO3a, GAPDH (used as a cytoplasmic marker), and Histone H3 (used as a nuclear marker). The virtual absence of the GAPDH signal in the nuclear fraction (N) and the Histone H3 signal in the cytoplasmic fraction (C) demonstrates the correct separation of the two cellular fractions. (B) Densitometric analysis of the FOXO3a signal. Data in panel B are expressed as mean ± SD and are representative of three independent experiments. **** indicates a significant difference of p < 0.0001 between the same cellular fractions in Fanc-A corr and Fanc-A scr. ### indicates a significant difference of p < 0.0001 between the same cellular fractions in Fanc-A + miR-29a-3p and Fanc-A scr.

### The miR-29a-3p-induced nuclear translocation of FOXO3a depends on the modulation of its hyperphosphorylation through AKT and SGK1 pathways

FOXO3a migration from cytoplasm to nucleus depends on different post-transcriptional modifications^36–38^, including the phosphorylation on Ser253 by AKT phosphorylated on Ser473, which is the principal post-translational modification that prevents FOXO3a entry into the nucleus^39,40^. Therefore, the phosphorylation levels of these two proteins were evaluated by WB analysis. Our results show that Fanc-A lymphoblasts displayed higher phosphorylation of Ser253 p-FOXO3a (Figure 5A and 5B) and Ser473 p-AKT (Figure 5A and 5C) compared to Fanc-A corrected cells, which was reverted by miR-29a-3p treatment. Furthermore, since SGK1 shares the same phosphorylation site on FOXO3a as AKT^41,42^ when phosphorylated on Ser422, the expression of phosphorylated and total forms of SGK1 in Fanc-A cells has been evaluated, observing a hyperphosphorylation in Fanc-A cells compared to Fanc-A corr, which was reduced after miR-29a-3p transfection(Figure 5A and 5D).

**Figure 5.**
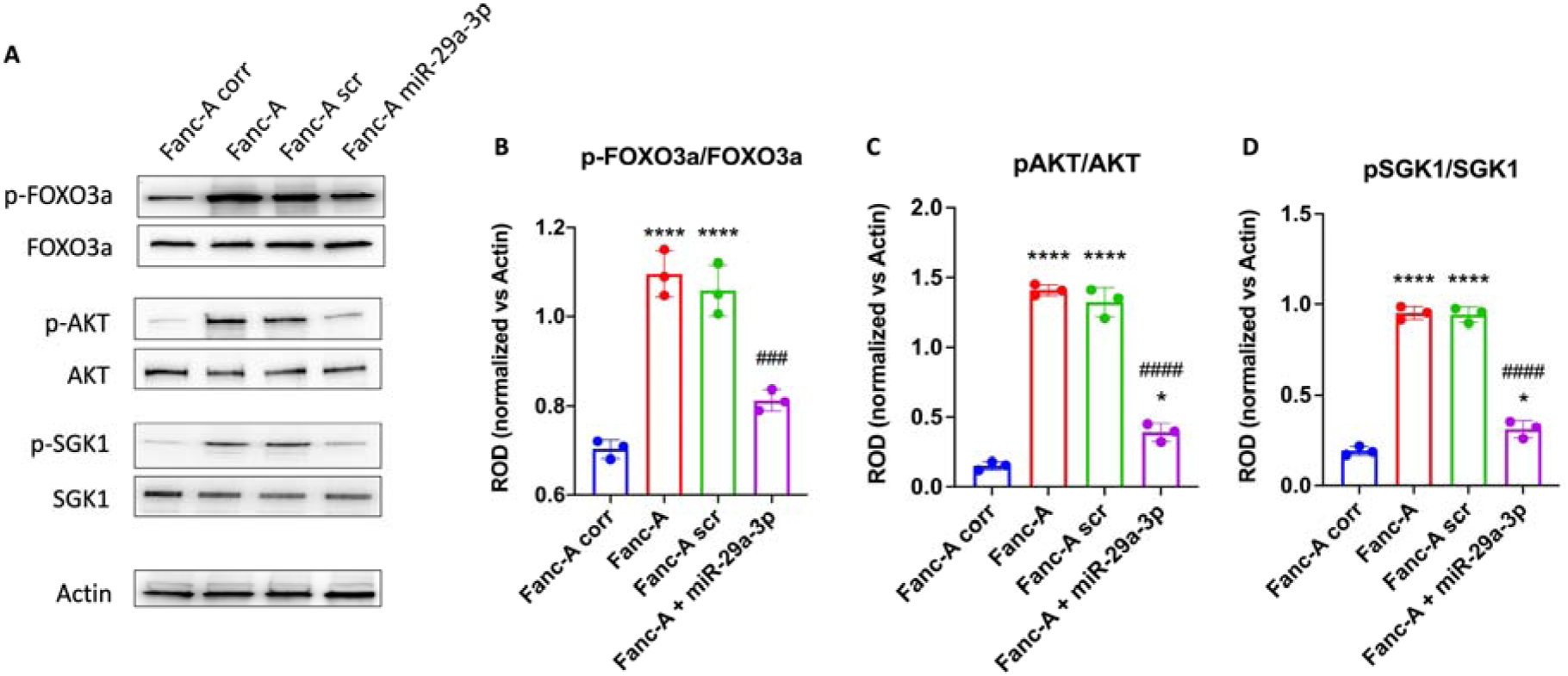
miR-29a-3p transfection modulates the FOXO3a,AKT, and SGK1 phosphorylation in Fanc-A lymphoblasts. All analyses were conducted on Fanc-A lymphoblasts corrected with the WT *Fanc-A* gene (Fanc-A corr), Fanc-A lymphoblasts (Fanc-A), Fanc-A lymphoblasts transfected with a miRNA mimic negative control for 48h (Fanc-A scr), and Fanc-A lymphoblasts transfected with miR-29a-3p for 48h (Fanc-A + miR-29a-3p). (A)Representative WB signals of:phospho-FOXO3a (Ser253); total FOXO3a; phospho-AKT (Ser473); total AKT; phospho-SGK1 (Ser422); total SGK1. Actin signal was used as housekeeping. (B) The ratio ofphosphorylated and total forms of FOXO3a signals. (C) The ratio ofphosphorylated and total forms of AKT signals. (D) The ratio of phosphorylated and total forms of SGK1 signals. Data in panels B, C, and D are expressed as mean ± SD and are representative of three independent experiments. * and**** indicate a significant difference for p <0.05 or 0.0001 between Fanc-A corr and other samples. ### and #### indicate a significant difference for p <0.001 or 0.0001, respectively, between Fanc-A + miR-29a-3p and Fanc-A scr.

### miR-29a-3p and TGF-β pathway modulate each other

Since the miR-29a-3p expression is negatively regulated by the TGF-β pathway hyperactivation^43^, which is a hallmark of FA cells^44^, its expression has been evaluated in the presence of Luspatercept, a TGF-β pathway inhibitor acting on SMAD2/3 signaling^45^. Data show that FA lymphoblast treated with Luspatercept increased the miR-29a-3p expression, reaching the level of control cells (Figure 6).

**Figure 6.**
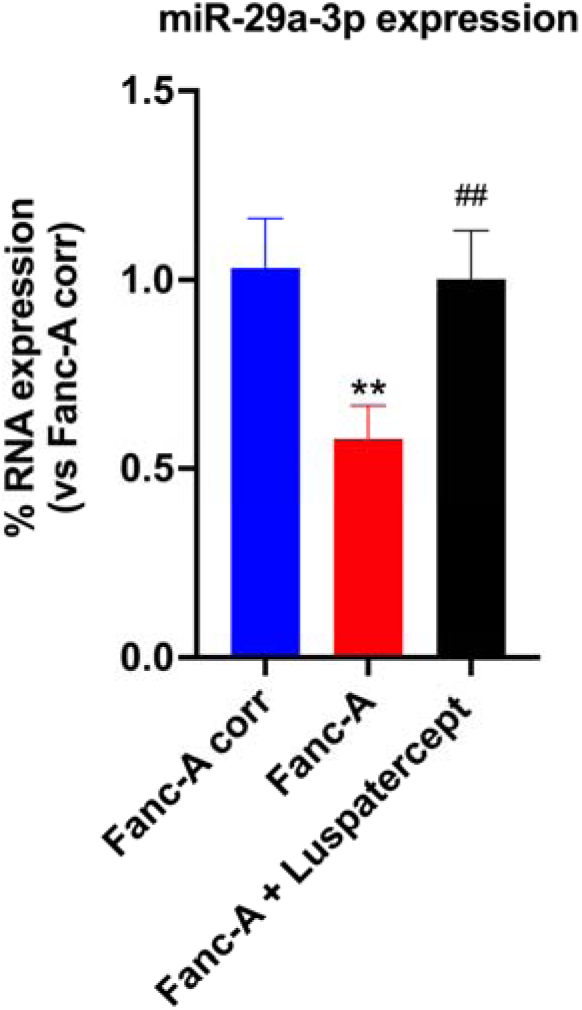
miR-29a-3p expression in Fanc-A lymphoblasts treated with Luspatercept. The graph shows the miR-29a-3p expression in Fanc-A lymphoblasts corrected with the WT *Fanc-A* gene (Fanc-A corr), Fanc-A lymphoblasts (Fanc-A), Fanc-A lymphoblasts treated with Luspatercept (TGF-beta pathway inhibitor) for 48h (Fanc-A + Luspatercept). Data are expressed as mean ± SD and are representative of three independent experiments **indicates a significant difference for p <0.01 between Fanc-A corr and Fanc-A. ## indicates a significant difference for p <0.01between Fanc-A and Fanc-A + Luspatercept.

On the other hand, miR-29a-3p also affects TGF-β signaling, as miR-29a-3p transfection in Fanc-A lymphoblasts reduced SMAD3 hyperphosphorylation to levels comparable to those observed in corrected cells (Figure 7).

**Figure 7.**
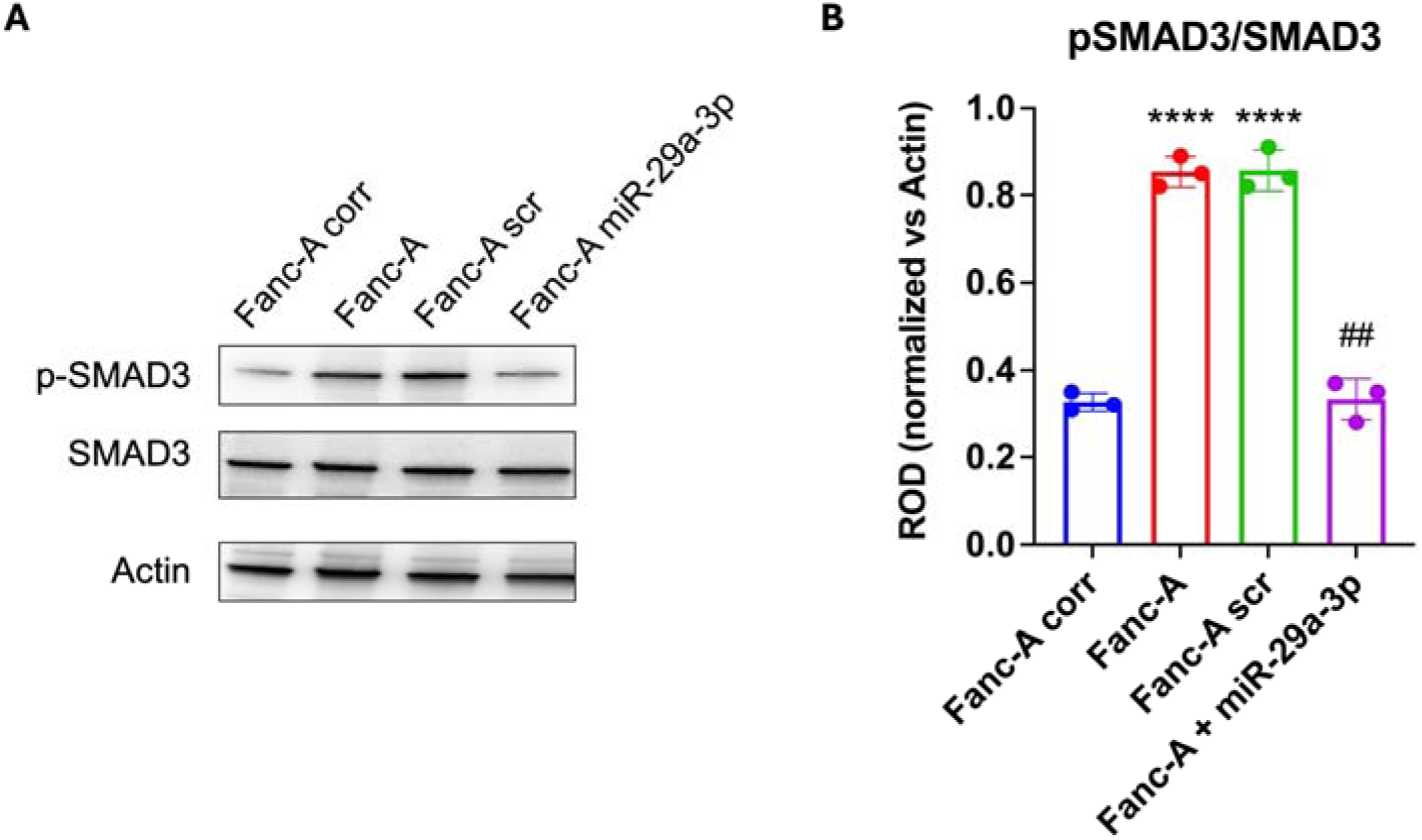
SMAD3 phosphorylation in Fanc-A lymphoblasts transfected with miR-29a-3p. All analyses were conducted on Fanc-A lymphoblasts corrected with the WT *Fanc-A* gene (Fanc-A corr), Fanc-A lymphoblasts (Fanc-A), Fanc-A lymphoblasts transfected with a miRNA mimic negative control for 48h (Fanc-A scr), and Fanc-A lymphoblasts transfected with miR-29a-3p for 48h (Fanc-A + miR-29a-3p). (A)Representative WB signals of phospho-SMAD3 (Ser423/425) and total SMAD3; Actin signal was used as housekeeping. (B) The ratio ofphosphorylated and total forms of SMAD3 signals. Data in panel B are expressed as mean ± SD and are representative of three independent experiments. **** indicates a significant difference for p<0.0001 between Fanc-A corr and other samples. ## indicates a significant difference for p < 0.01 between Fanc-A + miR-29a-3p and Fanc-A scr.

### Inhibition of TGF-βor IGF1 signaling reduces the oxidative damage and improves the oxidative phosphorylation in Fanc-A lymphoblasts

As shown in Figure 6, the TGF-βsignal reduction leads to a recoveryof miR-29a-3p content in FA lymphoblasts, suggesting the evaluation of possible effects of Luspatercept on antioxidant defenses, DNA damage, and energy metabolism. Moreover, since FA cells displayed elevated IGF1 expression compared to the control, which, however, decreased after transfection with miR-29a-3p (Figure 3), the same investigation was considered appropriate for Klotho, an IGF1 signaling inhibitor ^46^.

The data show that both the inhibition of the TGF-β pathway and IGF1 signaling exhibit the same trend observed after transfecting FA cells with miR-29a-3p. In particular,Luspatercept and Klothotreatmentsincreased AO response (Figure 8A), an evident reduction in oxidative stress accumulation(Figure 8B-C),a recovery of mitochondrial function(Figure 8D-G), and of cellular energy status (Figure 8H-L).

**Figure 8.**
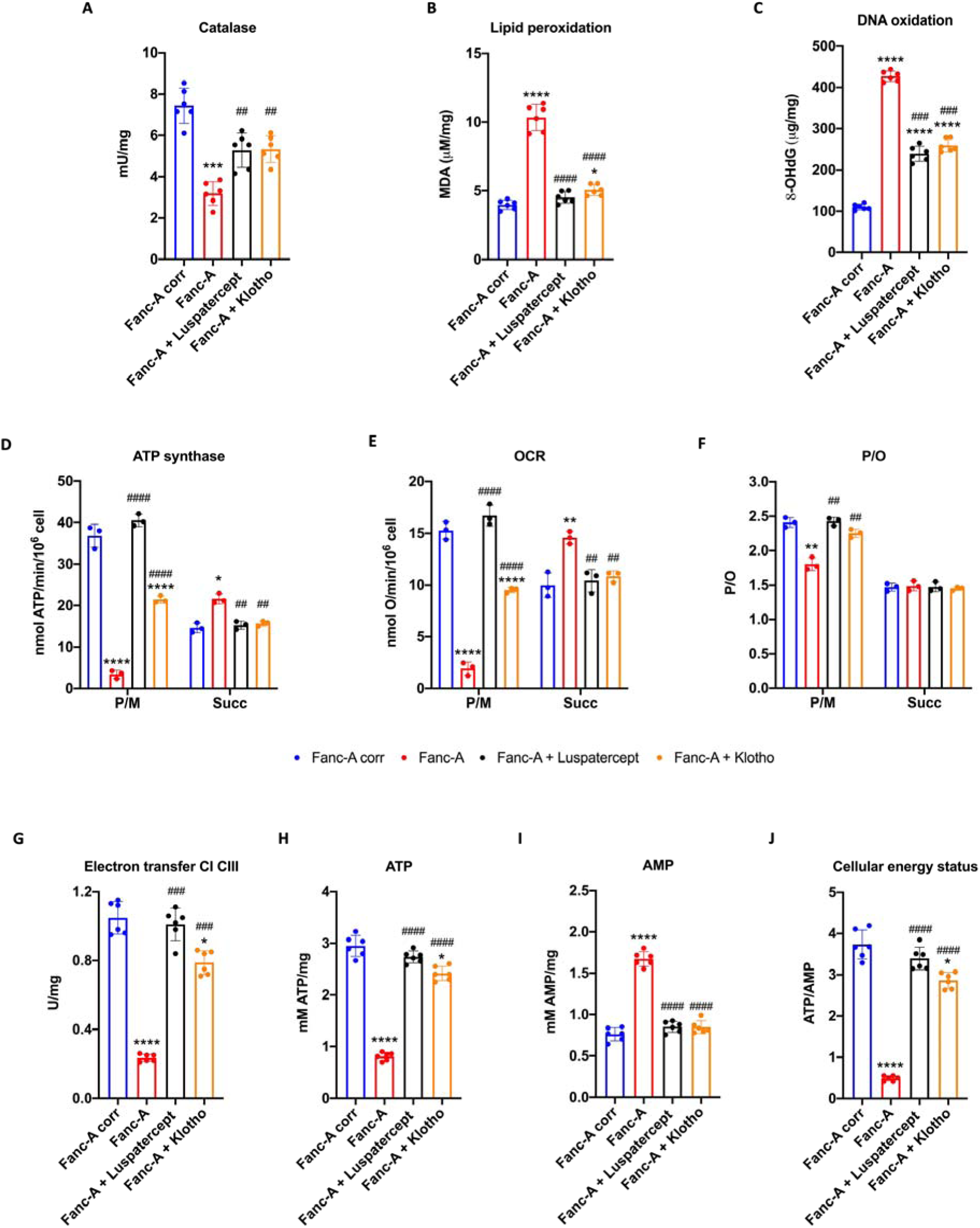
Antioxidant defenses, oxidative stress, and energy metabolism were modulated by Luspaterceptor Klothotreatment in Fanc-A lymphoblasts. All analyses were conducted on Fanc-A lymphoblasts corrected with the WT *Fanc-A* gene (Fanc-A corr), Fanc-A lymphoblasts (Fanc-A), Fanc-A lymphoblasts treated with Luspatercept (TGF-β pathway inhibitor) for 48h(Fanc-A + Luspatercept), and Fanc-A lymphoblasts treated with Klotho (IGF1 signaling inhibitor) for 48h (Fanc-A + Klotho). (A)Catalase activity as an antioxidant defense marker. (B) Malondialdehyde (MDA) intracellular concentration, as a lipid peroxidation marker. (C) 8-hydroxy-2’-deoxyguanosine (8-OHdG) content as a DNA oxidation marker. (D) ATP synthesis through F_o_F_1_-ATP synthase. (E) Oxygen consumption rate (OCR). (F) P/O value, an OxPhos efficiency marker. For Panels D, E, and F, the analyses were conducted in the presence of pyruvate plus malate (P/M) or succinate (Succ) to induce the OxPhos pathways led by Complex I or Complex II, respectively. (G) Electron transfer between Complexes I and III. (H) Intracellular ATP content. (I) Intracellular AMP content. (J) Cellular energy status is obtained by calculating the ATP/AMP ratio. Data are expressed as mean ± SD and are representative of three independent experiments for Panels A and E-G and six independent experiments for Panels B-D and H-K. *, **, ***, and **** indicate a significant difference for p <0.05, 0.01, 0.001, and 0.0001, respectively, between Fanc-A corr and Fanc-A. ##, ###, and ### indicate a significant difference for p < 0.01, 0.001, and 0.0001, respectively, between Fanc-A and Fanc-A + Luspatercept.

### Luspatercept and Klotho treatments reduce the hyperphosphorylation of FOXO3a, SGK1, and AKT

Since Luspatercept and Klotho treatments modulate miR-29a-3p expression and downstream function, respectively, their effects on the hyperphosphorylation of FOXO3a, AKT, and IGF1 have been investigated. Data reported in Figure 9 show that bothtreatments can reduce the phosphorylation level of miR-29a-3p targets, restoring levels similar to the control.

**Figure 9.**
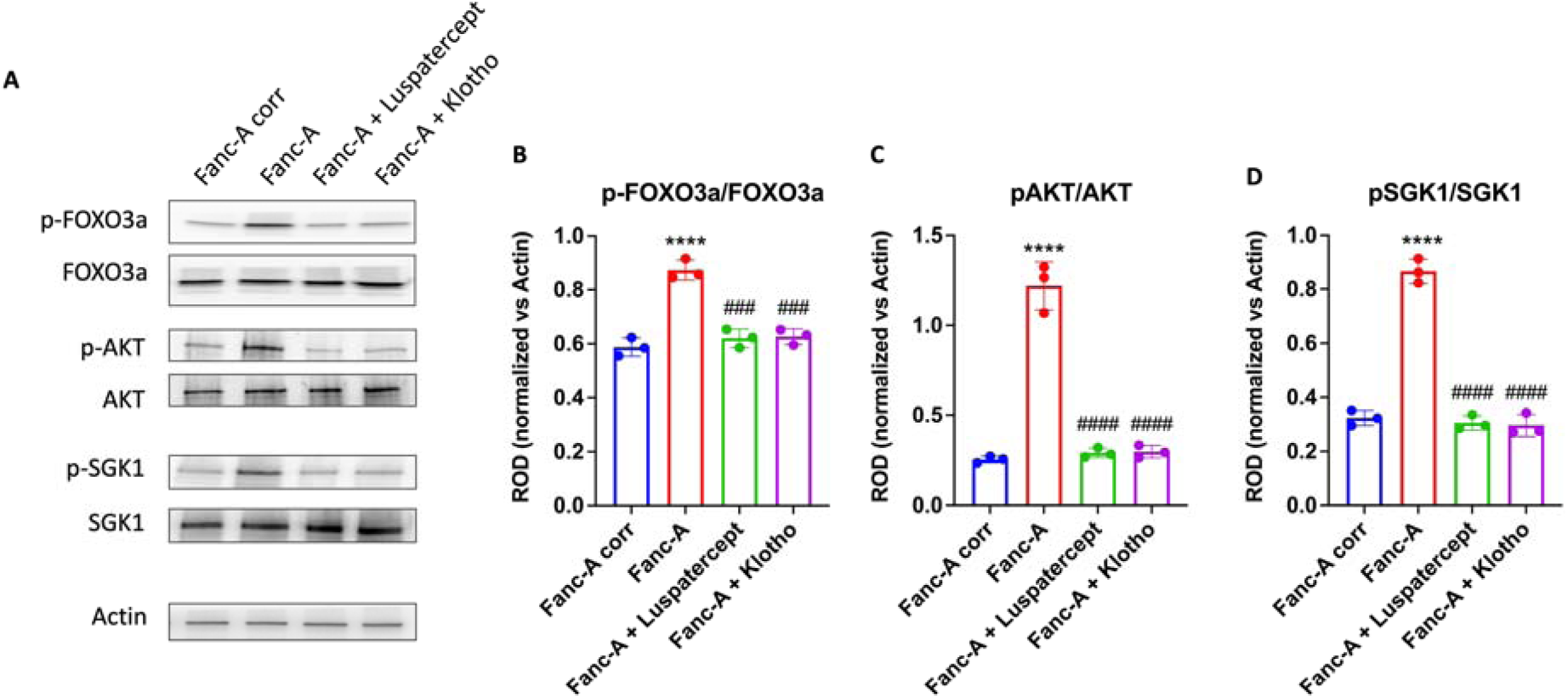
Luspatercept and Klothotreatmentsmodulate the FOXO3a, AKT, and SGK1 phosphorylation in Fanc-A lymphoblasts. All analyses were conducted on Fanc-A lymphoblasts corrected with the WT *Fanc-A* gene (Fanc-A corr), Fanc-A lymphoblasts (Fanc-A), Fanc-A lymphoblasts treated with Luspatercept (TGF-beta pathway inhibitor) for 48h(Fanc-A + Luspatercept), and Fanc-A lymphoblasts treated with Klotho (IGF1 signaling inhibitor) for 48h (Fanc-A + Klotho). (A) Representative WB signals of: phospho-FOXO3a (Ser253); total FOXO3a; phosphor-AKT (Ser473); total AKT; phosphor-SGK1 (Ser422); total SGK1. Actin signal was used as housekeeping. (B) The ratio of phosphorylated and total forms of FOXO3a signals. (C) The ratio of phosphorylated and total forms of AKT signals. (D) The ratio of phosphorylated and total forms of SGK1 signals. Data in panels B, C, and D are expressed as mean ± SD and are representative of three independent experiments. **** indicate a significant difference for p <0.0001 between Fanc-A corr and other samples. ### and #### indicate a significant difference for p <0.001 or 0.0001, respectively, between Fanc-A + miR-29a-3p and Fanc-A scr.

## DISCUSSION

The data presented herein provide novel insights into the role of the miR-29a-3p and TGFβ axis in FA pathogenesis, particularly concerning mitochondrial dysfunction, redox balance, and DNA damage accumulation.

These findings suggest that, although the defect in DNA damage repair is the principal cause of FA, additional molecular mechanisms contribute to maintaining an altered cellular state, perpetuating a vicious cycle that exacerbates DNA damage over time.

Previous studies have highlighted the dysregulation of several miRNAs in FA cells ^30,31^. However, to the best of our knowledge, this is the first study to demonstrate that the downregulation of miR-29a-3p impacts energetic and redox imbalances typical of FA cells. In detail, we report that the miR-29a-3p transfection improved OxPhos in terms of function and coupling and promoted the antioxidant enzyme expression and activity in FA lymphoblasts and fibroblasts. This enhancement of redox balance prevented membrane lipoperoxidation, as evidenced by reduced MDA levels, and mitigated DNA damage, as indicated by decreased 8-OHdG content and reduced H2AX phosphorylation. On the other hand, miR-29a-3p has emerged as a critical regulator, modulating the expression of genes involved in mitochondrial activity ^47^, redox balance^32^, DNA damage ^32^, and cell proliferation ^48^.

To understand the mechanism underlying the influence of miR-29a-3p on the energy metabolism and redox balance of FA cells, we focused our attention on FOXO3a, a miR-29a-3p-targeted transcription factor playing a pivotal role in the mitochondrial metabolism modulation^47^. On the other hand, miR-29a-3p appears essential for the self-renewal of hematopoietic stem cells just by controlling mitochondrial function^49^. Our data show that FA cells displayed a threefold higher expression of the FOXO3 gene compared to control cells. This elevated expression in FA cells could explain the metabolic alterations characterizing FA cells as FOXO3a activation causes a reduction in mitochondrial DNA copy number, mitochondrial protein expression, respiratory complexes, and mitochondrial respiratory activity^50^. This hypothesis is confirmed by the recovery of OxPhos activity and efficiency as well as the cellular energy status improvement observed after the reduction of FOXO3 gene expression following the transfection with miR-29a-3p both in FA lymphoblasts and fibroblasts. In addition, behind the FOXO3 gene expression, it is necessary to consider that FOXO3a protein functiondepends on its cellular localization, which is regulated by various post-translational modifications; for example, FOXO3a nuclear translocation is promoted by the macrophage stimulating 1 (MST1)-induced phosphorylation while it is inhibited by thephosphorylation on Thr32 and Ser253 by Ser473-phosphorylated AKT^41^. Since AKT hyperphosphorylation is a hallmark of FA cells ^51^, it is quite surprising to observe that, in FA lymphoblasts, FOXO3a appears hyperphosphorylated at Ser253 and accumulates in the cytoplasmic fraction. Interestingly, both FOXO3a and AKT hyperphosphorylation were reduced in cells transfected with miR-29a-3p, leading to an increase in FOXO3a levels in the nuclear fraction, suggesting that the AKT pathway is also regulated by miR-29a-3p^52^. Indeed, a recent study by Pang et al. proposes that the FOXO3a localization in nuclei depends on the mono-ubiquitination of FANCD2 and is independent of AKT phosphorylation. However, this discrepancy suggests that the regulation of FOXO3a nuclear localization is highly complex and involves additional post-translational modifications^53^.

Since FOXO3a also promotes ROS detoxification by increasingcellular antioxidant defense ^54^and genome stability ^35^, it is plausible to speculate that the miR-29a-3p-induced FOXO3 gene and protein modulation also leadsto the increase in catalase activity and the decrease of the DNA damage accumulation observed in transfected FA lymphoblasts and fibroblasts.

The SGK1, another miR-29a-3p target gene, also modulates FOXO3a phosphorylation and the consequent cellular localization. In detail, SGK1 phosphorylated on Ser422 leads to the FOXO3a negative modulation^41^. Regarding this, our data show that FA lymphoblasts are characterized by elevated SGK1 gene expression and hyperphosphorylation of the SGK1 protein, which return to levels similar to those of controls following transfection with miR-29a-3p. Therefore, these data suggest that miR-29a-3p transfection exerts a dual effect on FOXO3a: it acts as a direct modulator of its gene expression while also influencing its cellular localization and function through the regulation of AKT and SGK1, another target gene of miR-29a-3p.

To investigate the mechanism underlying the downregulation of miR-29a-3p in FA cells, we focused on the TGF-β signaling, as this pathway is hyperactivated in FA cells^55^. In addition, the miR-29a-3p expression is modulated by TGF-β through the signal transducer SMAD3 ^56^. Indeed, our data show that inhibition of TGF-β signaling following treatment with Luspatercept leads to an increased expression of miR-29a-3p in FA lymphoblasts, which, in turn, exerts positive effects on mitochondrial metabolism, cellular energy status, and the activation of antioxidant defenses. The same results were also observed in the presence of Klotho, an inhibitor of IGF1 signaling, which is an effector of the TGF-β pathway^57^. IGF1 expression is elevated in FA cells and is regulated by miR-29a-3p. This finding is particularly interesting given that FA patients are characterized by borderline hyperglycemia^58^, which could lead to increasing in IGF1 signaling.

The inhibition of the TGF-β signal throughLuspatercept or Klotho also causes a reduction of DNA damage accumulation, as demonstrated by the decrement in 8-OHdG content and H2AX phosphorylation. On the other hand, in FA cells, the hyperactivation of TGF-β signaling promotes DNA repair via non-homologous end joining (NHEJ), an error-prone repair pathway that contributes to toxicity in FA hematopoietic stem cells^55^. Conversely, inhibition of the TGF-β pathway through silencing of the SMAD3 gene in FA mice and human cells modulates the expression of DNA repair genes in favor of homologous recombination (HR) over NHEJ, resulting in increased growth of hematopoietic progenitors and rescue of bone marrow failure ^29^.

In addition, TGF-β signaling inhibition plays a pivotal role in modulating the PI3K/AKT pathway ^59,60^ also influencing FOXO3a, as demonstrated by the decreased phosphorylation of AKT, SGK1, and FOXO3a following treatment with Luspatercept and Klotho. Interestingly, miR-29a-3p is also a modulator of the TGF-β pathway, as its transfection into FA cells reduces the phosphorylation of SMAD3, a key effector of the TGF-β pathway. In other words, the data suggest that miR-29a-3p and TGF-β are connected through a negative feedback loop that mutually regulates their expression. Therefore, if increased miR-29a-3p expression can restore mitochondrial functionality and redox balance while reducing DNA damage accumulation in FA cells through the modulation of AKT, SGK1, and FOXO3a, it is plausible to hypothesize that thismodulation, in turn, reduces the release of pro-inflammatory cytokines, leading to a consequent decrease of the TGF-β signaling, further promoting miR-29a-3p expression.

## CONCLUSIONS

The data presented in this study demonstrate the central role of miR-29a-3p and the TGF-β pathway in the pathogenesis of FA, as the overexpression of the former and the reduction of the latter promote the recovery of metabolic functionality, the restoration of redox balance, and the reduction of DNA damage accumulation. Furthermore, the literature reports that miR-29a-3p plays a pivotal role in hematopoiesis, supporting self-renewal, lineage commitment, and HSC differentiation^49^. In addition, altered miR-29a-3p expression is associated with the development of Head and Neck Squamous Cell Carcinoma ^61^, a type of cancer highly prevalent in individuals with FA^62^.

These findings also suggest that the cellular dysfunctions characteristic of FA, which lead to DNA damage accumulation, depend on the activation of multiple, partially redundant signaling pathways. Consequently, any potential therapeutic approach should aim to modulate these pathways at multiple levels to ensure effectiveness.

## MATERIALS AND METHODS

### Cell Culture and treatments

Fanc-A lymphoblast cell lines and FANC-A primary fibroblast cell lines (Fibro FA) that carried out different mutations of the FANC-A gene were obtained from the ‘‘Cell Line and DNA Biobank from Patients affected by Genetic Diseases’’ (G. Gaslini Institute) - Telethon Genetic Biobank Network (Project No. GTB07001). As controls, isogenic FA-corr cell lines generated by the same FANC-A lymphoblast and fibroblast cell lines corrected with S11FAIN retrovirus (Lympho FA-corr and Fibro FA-corr) were employed ^34^.

The study was conducted following the Declaration of Helsinki and approved by the regional ethics committee, protocol JS002, register number 037-21/01/2019. All the subjects or their legal guardians gave written informed consent to the investigation.

For lymphoblast cell lines, RPMI-1640 medium (GIBCO, Billing, MT, USA) containing 10% fetal bovine serum (FBS, Euroclone, Milano, Italy), 100 U/mL penicillin, and 100 μg/mL streptomycin (Euroclone, Milano, Italy) was used, and the cells were grown at 37 °C with a 5% CO_2_^34^. Primary fibroblasts were cultured as a monolayer in DMEM high glucose with glutamax® (GIBCO, Billing, MT, USA), containing 10% FBS (Euroclone, Milano, Italy), 100 U/mL penicillin, and 100 μg/mL streptomycin (Euroclone, Milano, Italy) at 37 °C with a 5% CO_2_^34^.

### FA lymphoblast and fibroblast cells transfection with miR-29a-3p

FA lymphoblast cells were transfected with miR-29a-3p mimic (ThermoFisher Scientific, Waltham, MA, USA), using the lipofectamineRNAiMAXTransfectionReagent(Invitrogen, Waltham, MA, USA) according to the manufacturer’s instruction. In detail, for subsequent RNA extraction, 500.000 cells grown in 6-well plates were transfected with 25 pmol of miR-29a-3p mimic or negative control mimic using 7.5μl lipofectamine. After 48h, cells were harvested, washed with PBS, lysed, and RNA was extracted.For the subsequent biochemical analysis, 7.5×10^6^ cells grown in 75 cm^2^ flasks were transfected with 187.5 pmol of miR-29a-3p mimic or negative control mimic using 56.25 μl lipofectamine. Cells were processed 48h post-transfection.

FA fibroblast cells were transfected with miR-29a-3p mimic (Thermo Fisher Scientific, Waltham, MA, USA) using the same transfectionreagent(Invitrogen, Waltham, MA, USA). In detail, for the subsequent RNA extraction, 75.000 cells grown in 6-well plates were transfected with 25 pmol of miR-29a-3p mimic or negative control mimic using 7.5 μl lipofectamine. After 48h, cells were harvested, washed with PBS, lysed, and RNA was extracted.For subsequent biochemical analysis, 500.000 cells grown in 175 cm^2^ flasks were transfected with 437.5 pmol of miR-29a-3p mimic or negative control mimic using 131,25 μllipofectamine. Cells were processed 48h post-transfection.

### FA lymphoblast cell treatments

To inhibit the TGF-β pathway, lymphoblasts were treated with Luspatercept, an inhibitor acting on the SMAD2/3 signaling ^45^. In detail, 500.000 cells were plated in a 2 ml culture medium in the presence of 10 μg/ml Luspatercept for 48h.

To inhibit IGF1 signaling, 500.000 lymphoblasts were treated with 400 pM Klotho ^46^ for 48h.

### In silico selection of pathway-related miR-29a-3p target genes

To identify putative miR-29a-3p target genes potentially involved in Fanconi cell metabolism impairment, we utilized the miRPathDB v2.0 database (https://mpd.bioinf.uni-sb.de). A curated list of miR-29a-3p-regulated genes associated with DNA damage response, oxidative stress, mitochondrial metabolism, lipid metabolism, and apoptosis was generated by evaluating the strength of predicted miRNA-target gene interactions using TargetScan (https://mpd.bioinf.uni-sb.de). Further refinement was conducted by assessing the role and the subcellular localization of each gene with the help of the NCBI Gene database (https://www.ncbi.nlm.nih.gov/gene) and GeneCards (https://www.genecards.org). The final list is presented in Table 1.

### RNA isolation to evaluate the expression of miR-29a-3p and FOXO3, SGK1, and IGF1 genes

RNA, including the small RNA fraction, was extracted using the RNeasy Plus Mini Kit (Qiagen, Hilden, Germany) according to the manufacturer’s protocol. The expression of miR-29a-3p was assessed using a specific TaqMan MicroRNA Assay (Applied Biosystems, Waltham, MA, USA). Briefly, 10 ng of RNA was reverse-transcribed using the TaqMan MicroRNA Reverse Transcription Kit and the specific RT primer. Real-time PCR was performed in triplicate with specific primers. miRNA expression levels were normalized to RNU44 expression.

To evaluate the expression of the FOXO3, SGK1, and IGF1 genes, 100 ng of RNA were reverse transcribed using the SuperScript VILO IV cDNA Synthesis Kit (Invitrogen, Waltham, MA, USA). The resulting cDNA was used for real-time PCR with primers provided in the specific TaqMan Gene Expression Assays (Applied Biosystems, Waltham, MA, USA). Gene expression levels were normalized to GAPDH expression. All experiments were performed in triplicate.

### Catalase activity evaluation

Catalase (CAT) activity was assayed as a marker of cellular antioxidant defenses. For each spectrophotometric assay, 50 μg of total proteins were used, following the H_2_O_2_ decomposition at 240 nm. The assay mix contained 50 mM phosphate buffer (pH 7.0) and 5 mM H_2_O_2_ (Sigma-Aldrich, St. Louis, MO, USA) ^21^.

### Oxidative stress markers evaluation

The thiobarbituric acid reactive substances (TBARS) assay was employed to quantify malondialdehyde (MDA), an indicator of lipid peroxidation. The TBARS reagent was prepared using 0.25 M HCl, 0.25 mM trichloroacetic acid, and 26 mM thiobarbituric acid (all sourced from Merck, Darmstadt, Germany). A total of 50 µg of protein, dissolved in 300 µl of Milli-Q water, was mixed with 600 µl of the TBARS solution. The reaction mixture was incubated at 95 °C for 1 hour and the absorbance was measured spectrophotometrically at 532 nm. Standard solutions of MDA, with concentrations ranging from 1 to 20 µM, were used to generate a calibration curve ^21^.

To measure 8-hydroxy-2-deoxyguanosine (8-OHdG), a biomarker of oxidative DNA damage, an ELISA kit (#ab201734, Abcam, Cambridge, UK) was utilized according to the manufacturer’s instructions.

### Evaluation of aerobic metabolism function and efficiency

The OxPhos function was assessed by measuring the oxygen consumption rate (OCR) and F_o_F_1_-ATP synthase activity.

OCR was determined using an amperometric electrode (Unisense Microrespiration, Aarhus, Denmark) in a closed chamber. For each test, 10^5^ cells were permeabilized for 1 minute with 0.03 mg/ml digitonin and then used in the assay. To activate the respiratory pathway led by Complex I or Complex II, 10 mM pyruvate with 5 mM malate (Merck, Darmstadt, Germany) or 20 mM succinate Merck, Darmstadt, Germany) were added, respectively ^21^.

F_o_F_1_-ATP synthase activity was measured in 10^5^ cells suspended in PBS plus 0.6 mM ouabain Merck, Darmstadt, Germany) and 0.25 mM di(adenosine)-5-penta-phosphate (an adenylate kinase inhibitor, Merck, Darmstadt, Germany). After 10 minutes of incubation, 10 mM pyruvate with 5 mM malate (Merck, Darmstadt, Germany) or 20 mM succinate (Merck, Darmstadt, Germany) were added to stimulate pathways mediated by Complex I or II, respectively. ATP production was quantified using a luminometer (GloMax® 20/20 Luminometer, Promega Italia, Milan, Italy),employing the luciferin/luciferase chemiluminescent method (ATP bioluminescence assay kit CLS II, #11699695001 Roche, Basel, Switzerland). Measurements were taken at 30-second intervals over 2minutes ^21^.

The OxPhos efficiency was determined by calculating the P/O ratio, which represents the amount of ATP synthesized aerobically per oxygen molecule consumed. Mitochondria with optimal efficiency exhibit P/O ratios around 2.5 for pyruvate and malate or 1.5 for succinate. Ratios below these thresholds suggest incomplete oxygen utilization for ATP production, potentially reflecting increased ROS generation ^21^.

### Assessment of ATP and AMP intracellular concentration and the consequent cellular energy status

ATP and AMP concentrations were assayed in 50 μg of total protein. ATP content spectrophotometric analysis was performed following the NADPreduction at 340 nm. The assay solution contained 100 mMTris-HCl (pH 8.0; Merck, Darmstadt, Germany), 0.2 mM NADP(Merck, Darmstadt, Germany), 5 mM MgCl_2_(Merck, Darmstadt, Germany), 50 mM glucose (Merck, Darmstadt, Germany), and 3 μg of pure hexokinase and glucose-6-phosphate dehydrogenase (Merck, Darmstadt, Germany)^20^.

AMP was measured spectrophotometrically following the NADH oxidation at 340 nm. The reaction medium was composed of 100 mMTris-HCl (pH 8.0; Merck, Darmstadt, Germany), 5 mM MgCl_2_ (Merck, Darmstadt, Germany), 0.2 mM ATP (Merck, Darmstadt, Germany), 10 mMphosphoenolpyruvate (Merck, Darmstadt, Germany), 0.15 mM NADH (Merck, Darmstadt, Germany), 10 IU adenylate kinase, 25 IU pyruvate kinase, and 15 IU lactate dehydrogenase (Merck, Darmstadt, Germany) ^20^.

The cellular energy status was calculated as the ratio between intracellular concentration of ATP and AMP (ATP/AMP ratio)^20^.

### Evaluation of electron transfer between Complex I and Complex III

To evaluate the electron transfer between Complex I and Complex III, a spectrophotometric assay was performed following the reduction of oxidized cytochrome *c* (cyt*c*) at 550 nm in the presence of NADH. The assay medium contained 50 mMTris-HCl (pH 7.4; Merck, Darmstadt, Germany), 5 mMKCl (Merck, Darmstadt, Germany), 2 mM MgCl_2_ (Merck, Darmstadt, Germany), 0.5 M NaCl (Merck, Darmstadt, Germany), 0.03% oxidized cyt*c*(Merck, Darmstadt, Germany), and 0.6 mM NADH (Merck, Darmstadt, Germany) ^20^.

### Cellular fraction separation

To obtain nuclei-enriched fractions (N), cells were homogenated in a 0.25 M sucrose solution. This homogenate (H) was centrifuged at 800 g for 10 min. The supernatant was collected and used as a cytoplasmic fraction (C), and the pellet was resuspended in 0.25 M sucrose solution and centrifuged again at 800 g for 10 min to obtain N.

### Western Blot Analysis

Denaturing electrophoresis (SDS-PAGE) was performed on 30 μg of proteins employing a 4–20% gradient gel (BioRad, Hercules, CA, USA). The following primary antibodies were used: phospho-H2AX (#05-636, Merck, Darmstadt, Germany). phospho-FOXO3a (#SAB5701786 Merck, Darmstadt, Germany), FOXO3a (#2497S, Cell Signaling Technology, Beverly, MA, USA), GAPDH (#2118, Cell Signaling Technology, Beverly, MA, USA), H3 (#4499, Cell Signaling Technology, Beverly, MA, USA), phospho-AKT (#4060S, Cell Signaling Technology, Beverly, MA, USA), AKT (#4691S, Cell Signaling Technology, Beverly, MA, USA), phospho-SGK1 (#5599, Cell Signaling Technology, Beverly, MA, USA), SGK1 (#711183, Thermo Fisher Scientific, Waltham, MA, USA), phospho-SMAD3 (#9520S, Cell Signaling Technology, Beverly, MA, USA), SMAD3 (#9523S, Cell Signaling Technology, Beverly, MA, USA), and β-Actin (#MA1-140, ThermoFisher Scientific, Waltham. MA, USA). All primary antibodies were diluted following the manufacturer’s instructions in PBS plus 0.15% Tween 20 (PBSt; Roche, Basel, Switzerland). Specific secondary antibodies were employed (Merck, Darmstadt, Germany, all diluted 1:10,000 in PBSt). Bands were detected in the presence of an enhanced chemiluminescence substrate (ECL, BioRad, Hercules, CA, USA), bya chemiluminescence system (Alliance 6.7 WL 20M, UVITEC, Cambridge, UK). Band intensity was evaluated by UV1D software (UVITEC, Cambridge, UK). All bands of interest were normalized versus the actin signal detected on the same membrane.

### Statistical Analysis

Data were analyzed using one-way ANOVA followed by Tukey’s multiple comparison test by Prism 9 Software (GraphPad Software Inc., Boston, MA, USA). Data are expressed as mean ± SD and are representative of at least three independent experiments. An error with a probability of p<0.05 was considered significant.

## ACKNOWLEDGEMENTS

We want to acknowledge ERG SpA, Cambiaso and Risso, Rimorchiatori Riu-niti, and Saar Depositi Oleari Portuali for supporting the activity of the Clinical and Experimental Hematology Unit of the G. Gaslini Institute.

## AUTHOR CONTRIBUTIONS

Conceptualization, S.R., E.C., P.D., and Si.R.; investigation, N.B., S.R., V.C., M.B., M.S:, F.C., E.C., and Si.R.; resources, C.D., E.C., and Si.R.; data curation, S.R., E.C., Si.R.; writing, N.B., S.R., V.C., P.D., E.C., Si.R; supervision, C.B., F.P., C.D.

All authors have read and agreed to the published version of the manuscript.

## DATA AVAILABILITY

All data supporting the conclusions of this study can be found in the Article and Supplementary Material.

## SUPPLEMENTARY MATERIAL

**Figure S1.**
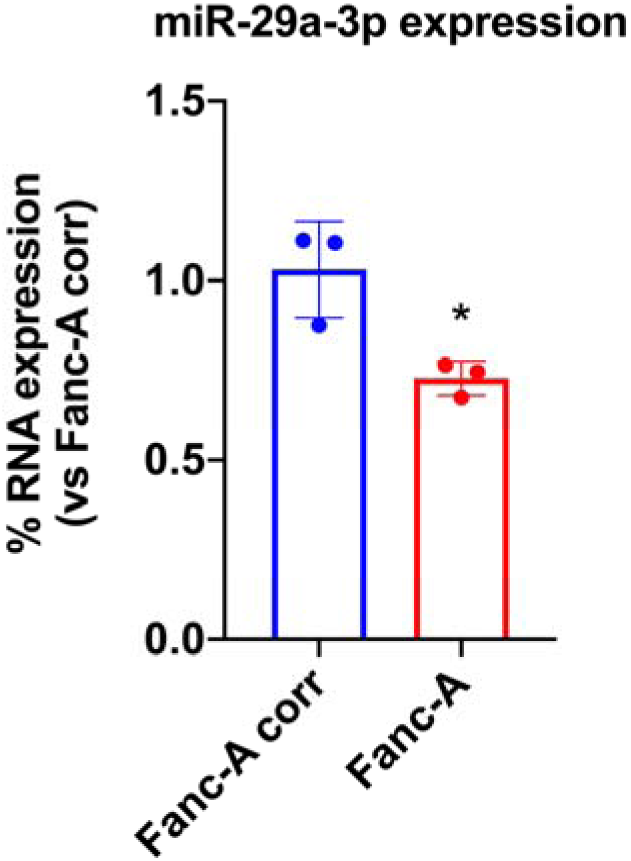
miR-29a-3p expression in Fanc-A fibroblasts. The graph shows the comparison of miR-29a-3p expression between Fanc-A fibroblasts corrected with the WT *Fanc-A* gene (Fanc-A corr) and Fanc-A fibroblasts (Fanc-A). RNU44 was used as a reference control.Data are expressed as mean ± SD and are representative of three independent experiments. *indicates a significant difference for p < 0.05 between Fanc-A corr and Fanc-A.

**Figure S2.**
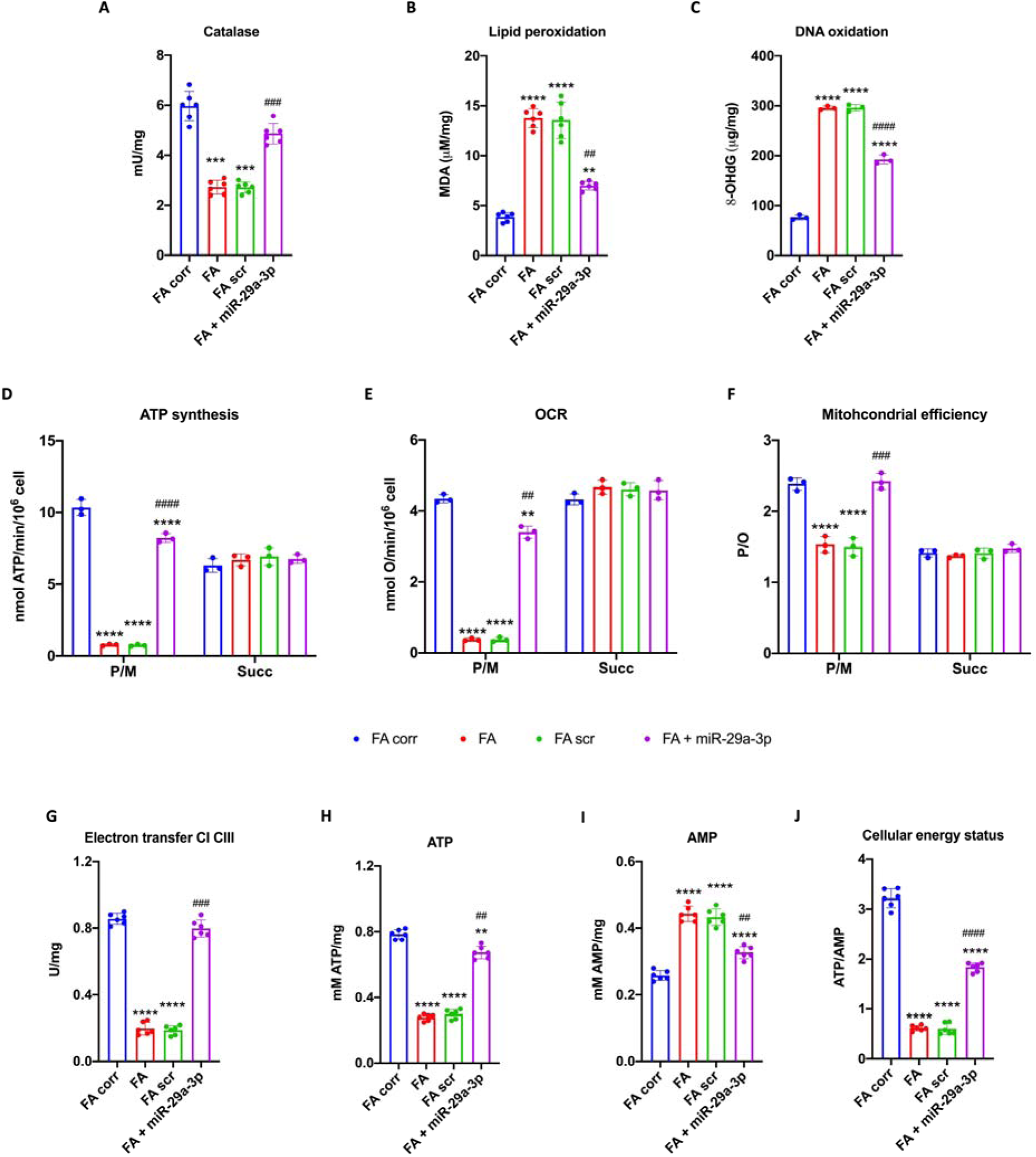
Antioxidant defenses, oxidative stress, and energy metabolism were modulated by miR-29a-3p expression in Fanc-A fibroblasts. All analyses were conducted on Fanc-A fibroblasts corrected with the WT *Fanc-A* gene (Fanc-A corr), Fanc-A fibroblasts (Fanc-A), Fanc-A fibroblasts transfected with empty vector for 48h (Fanc-A scr), and Fanc-A fibroblasts transfected with miR-29a-3p for 48h (Fanc-A + miR-29a-3p). (A) Catalase activity, as an antioxidant defenses marker. (B) Malondialdehyde (MDA) intracellular concentration, as a lipid peroxidation marker. (C) 8-hydroxy-2’-deoxyguanosine (8-OHdG) content, as a DNA oxidation marker. (D) ATP synthesis through F_o_F1-ATP synthase. (E) Oxygen consumption rate (OCR). (F) P/O value, an OxPhos efficiency marker. For Panels D, E, and F, the analyses were conducted in the presence of pyruvate plus malate (P/M) or succinate (Succ) to induce the OxPhos pathways led by Complex I or Complex II, respectively. (G) Electron transfer between Complexes I and III. (H) Intracellular ATP content. (I) Intracellular AMP content. (J) Cellular energy status is obtained by calculating the ATP/AMP ratio. Data are expressed as mean ± SD and are representative of three independent experiments for Panels D-F and six independent experiments for Panels A-C and G-J. **, ***, and **** indicate a significant difference for p < 0.01, 0.001, or 0.0001, respectively, between Fanc-A corr and Fanc-A or Fanc-A scr. ##, ###, and ### indicate a significant difference for p < 0.01, 0.001, or 0.0001, respectively, between Fanc-A + miR-29a-3p and Fanc-A or Fanc-A scr.

**Figure S3.**
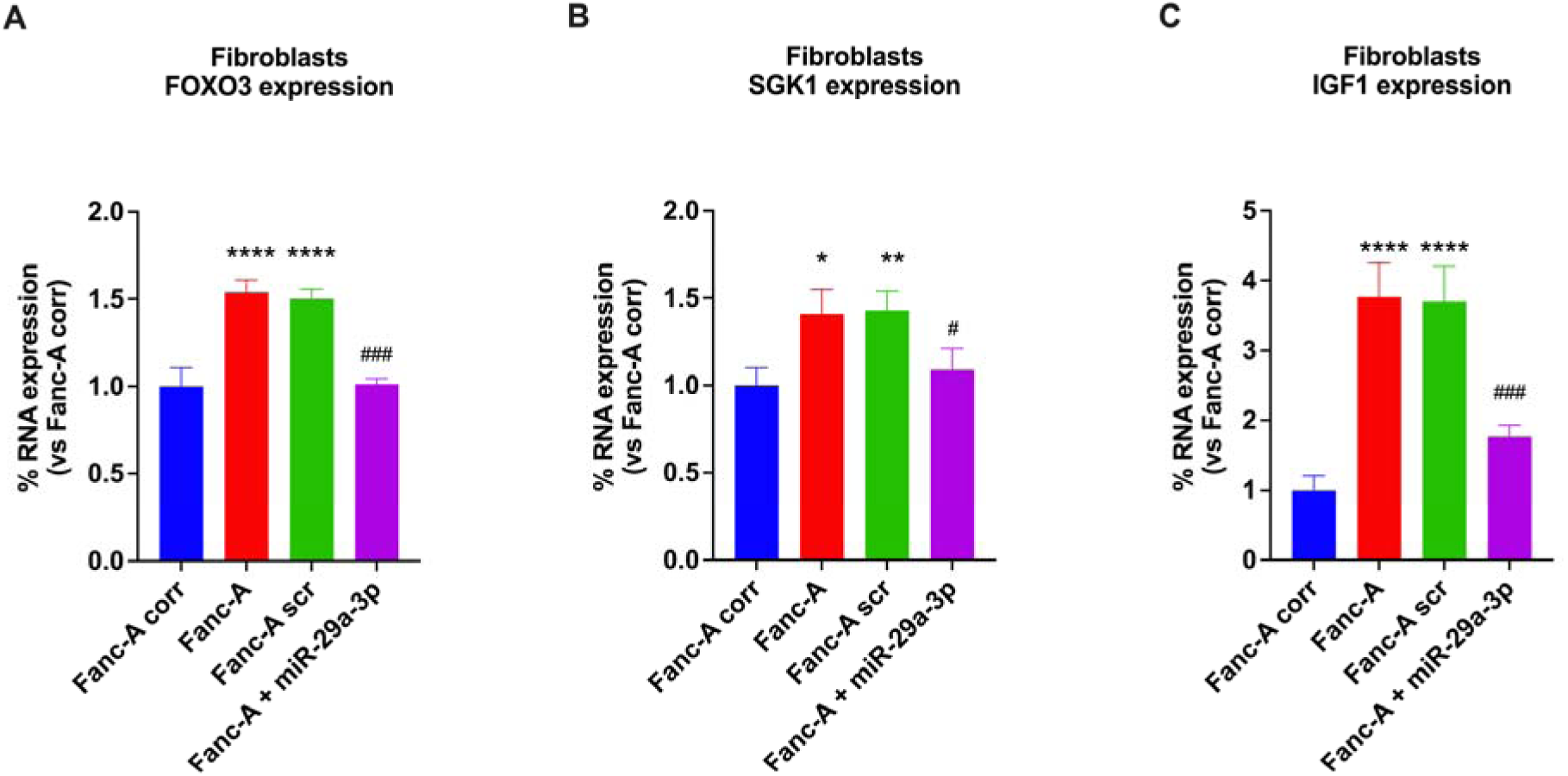
FOXO3, SGK1, and IGF1 expression in Fanc-A fibroblasts. Graphs show the comparison of FOXO3 (A), SGK1 (B), and IGF1 (C) expression in (i) Fanc-A cells corrected with the wt Fanc-A gene (Fanc-A corr), (ii) Fanc-A cells (Fanc-A), (iii) Fanc-A cells transfected with a miRNA mimic negative control for 48h (Fanc-A scr), and (iv) Fanc-A cells transfected with miR-29a-3p for 48h (Fanc-A + miR-29a-3p). GAPDH was used as the reference control. Data are expressed as mean ± SD and are representative of three independent experiments. *, ** or **** indicate a significant difference for p < 0.05, 0.01, or 0.0001, respectively, between Fanc-A corr and the other samples. # and ### indicates a significant difference for p < 0.05 or 0.001, respectively, between Fanc-A + miR-29a-3p and Fanc-A or Fanc-A scr.

